# The expected loss of feature diversity (versus phylogenetic diversity) following rapid extinction at the present

**DOI:** 10.1101/2022.10.09.511499

**Authors:** Marcus Overwater, Daniel Pelletier, Mike Steel

## Abstract

The current rapid extinction of species leads not only to their loss but also the disappearance of the unique features they harbour, which have evolved along the branches of the underlying evolutionary tree. One proxy for estimating the feature diversity (*FD*) of a set *S* of species at the tips of a tree is ‘phylogenetic diversity’ (*PD*): the sum of the branch lengths of the subtree connecting the species in *S*. For a phylogenetic tree that evolves under a standard birth–death process, and which is then subject to a sudden extinction event at the present (the simple ‘field of bullets’ model with a survival probability of *s* per species) the proportion of the original *PD* that is retained after extinction at the present is known to converge quickly to a particular concave function *φ*_*PD*_(*s*) as *t* grows. To investigate how the loss of *FD* mirrors the loss of *PD* for a birth–death tree, we model *FD* by assuming that distinct discrete features arise randomly and independently along the branches of the tree at rate *r* and are lost at a constant rate *v*. We derive an exact mathematical expression for the ratio *φ*_*FD*_(*s*) of the two expected feature diversities (prior to and following an extinction event at the present) as *t* becomes large. We find that although *φ*_*FD*_ has a similar behaviour to *φ*_*PD*_ (and coincides with it for *v* = 0), when *v >* 0, *φ*_*FD*_(*s*) is described by a function that is different from *φ*_*PD*_(*s*). We also derive an exact expression for the expected number of features that are present in precisely *one* extant species. Our paper begins by establishing some generic properties of FD in a more general (non-phylogenetic) setting and applies this to fixed trees, before considering the setting of random (birth–death) trees.

## 1 Introduction

Phylogenetic trees provide a way to quantify biodiversity and the extent to which it might be lost in the face of the current mass extinction event. One such biodiversity measure is *phylogenetic diversity* (PD), which associates to each subset *S* of extant species, the sum of the branch lengths of the underlying evolutionary tree that connects (just) these species to the root of the tree. This measure, pioneered by Daniel P. Faith [2], provides a more complete measure of biodiversity than merely counting the number of species in *S* (see e.g. [13]). Moreover, if new features evolve along the branches of a tree and are never lost, then the resulting features present amongst the species in the subset *S* (the *feature diversity* (FD) of *S*) are directly correlated with the phylogenetic diversity of *S* [25]. However, when features are lost, it has recently been shown mathematically that under simple (deterministic or stochastic) models of feature gain and loss, FD necessarily deviates from PD except for very trivial types of phylogenetic trees (Theorems 2 and 3 of [18]). More generally, the question of the extent to which PD captures feature or functional diversity has been the subject of considerable debate in the biological literature (see [1, 9, 10, 11, 16, 23, 22]).

In this paper, we investigate a related question: under a standard phylogenetic diversification model and a simple stochastic process of feature gain and loss, what proportion of feature diversity is expected to be lost in a mass extinction event at the present? And how does this ratio compare with the expected phylogenetic diversity that will be lost? Although the latter ratio (for PD) has been determined in earlier work, here we provide a corresponding result for FD, and show how it differs from PD when the rate of feature loss is non-zero. Our results suggest that the relative loss of FD under a mass extinction event at the present is greater than the relative loss of PD. We also investigate the number of features that are expected to be found in just one species at the present.

We begin by considering the properties of FD in a more general setting based purely on the species themselves (i.e. not involving any underlying phylogenetic tree or network) and then consider FD on fixed phylogenetic trees before presenting the results for (random) birth–death trees. Some of the results of these earlier sections are applied in the later sections.

## 2 General Properties of Feature Diversity without reference to phylogenies

This section considers the generic properties of expected FD for sets of species, and thus, no underlying phylogenetic tree or stochastic process that generates a tree, or model of feature evolution is assumed. We mostly follow the notation from [25].

**Definitions**

Let *X* be a labelled set of species, and for each *x* ∈ *X*, let *F*_*x*_ be a non-empty set of discrete features (e.g. genes, genomic inserts, traits) that are associated with species *x*.

- The collection of ordered pairs 𝔽 = {(*x, F*_*x*_) : *x* ∈ *X*} is called a *feature assignment* on *X*.
- For any subset *A* of *X*, let *ℱ*(*A*) = U_*x*∈*A*_ *F*_*x*_ and let *μ* : *ℱ*(*X*) *→* ℝ^*>*0^ be a function assigning some richness or novelty to a feature *f* ∈ *ℱ* (*X*).
- The *feature diversity* of some subset *A* of a set of species *X* is defined as,

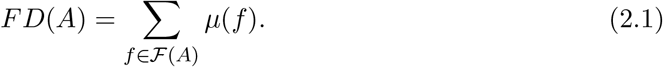

A default option for *μ* is to set *μ*(*f*) = 1, which simply counts the number of features present. Notice that for any subset *A* of *X* we have 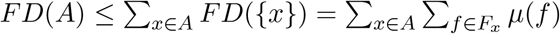, with equality if and only if no features are shared by any two species.

### 2.1 Feature diversity loss under a ‘Field of Bullets’ model of extinction at the present

#### Definition 2.1

Consider a sudden extinction event taking place across a set of species ([17]). In the *generalised field of bullets* (g-FOB) model, each species *x* ∈ *X* either survives the extinction event (with robability *s*_*x*_) or disappears (with probability 1 *−s*_*x*_), and these survival events are assumed to be independent among the species. We write **s**(*X*) (or, more briefly, **s**) to denote the vector (*s*_*x*_ : *x* ∈ *X*) and we let *χ* denote the (random) subset of *X* corresponding to the species that survive the extinction event. If *s*_*x*_ = *s* for all *x* ∈ *X*, then we have the simpler *field of bullets* (FOB) model.

We define the following quantity:

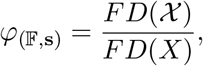

which is the proportion of feature diversity that survives the extinction event. In the case of the FOB model, we denote this ratio by *φ*_(𝔽,*s*)_.

#### Proposition 2.1

For the g-FOB model,

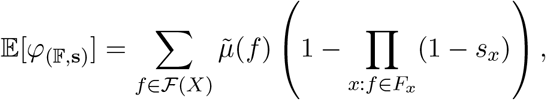

where 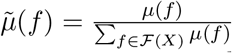 are the normalised *μ* values (which sum to 1).

For the FOB model, the equation in Proposition 2.1 simplifies to

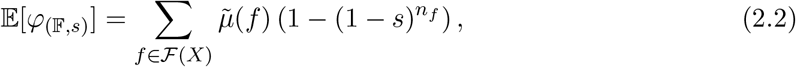

where *n*_*f*_ = |{*f* : ∃*x* : *f* ∈ *F*_*x*_}|, the number of species in *X* that possess feature *f*.

*Proof*. We have 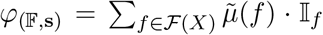 where 𝕀_*f*_ is the Bernoulli variable that takes the value 1 if at least one species in *X* with feature *f* survives the g-FOB extinction event, and 0 otherwise. Applying linearity of expectation and noting that 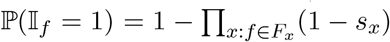 gives the result. □

Notice that 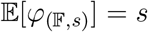 at *s* = 0, 1. The behaviour between these two extreme values of *s* is described next.

#### Proposition 2.2

Under the FOB model, the following hold:

i. 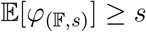 for all *s* ∈ [0, 1].
ii. 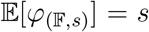 for a value of *s* ∈ (0, 1) if and only if the sets *F*_*x*_ in 𝔽 are pairwise disjoint, in which case 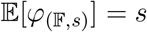 for all *s* ∈ [0, 1].
iii. If the sets *F*_*x*_ are not pairwise disjoint, then 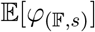 is a strictly concave increasing function of *s*.

*Proof*. Part (i): Since *n*_*f*_ *≥* 1 in Eqn. (2.2) and 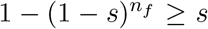 for all *n*_*f*_ *≥* 1, the claimed inequality is immediate. Part (ii): If *n*_*f*_ = 1 then 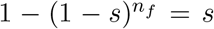 for all *s* ∈ [0, 1], and if *n*_*f*_ *>* 1 then 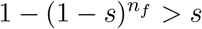 for every *s* ∈ (0, 1). Part (iii): By Eqn. (2.2),

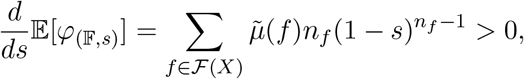

and

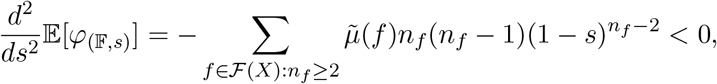

for all *s* ∈ (0, 1), from which the claimed results follow.

□

### 2.2 Approximating *φ*_(𝔽,*s*)_ by its expected value

For each *n ≥* 1, let *X*_*n*_ be a labelled set of *n* species with feature assignment 𝔽_*n*_. Let *χ*_*n*_ denote the (random) set of species after a g-FOB extinction event with survival probability vector **s**(*X*_*n*_), which assigns each *x* ∈ *X*_*n*_ a corresponding survival probability *s*_*n*_(*x*). Note that we make no assumption regarding how the species in *X*_*n*_ and *X*_*m*_ are related (e.g. they may be disjoint, overlapping or nested), or any apriori relationship between **s**(*X*_*n*_) and **s**(*X*_*m*_).

In the following result, we provide a sufficient condition under which *φ*_(𝔽*n*_,**s**_*n*_) is likely to be close to its expected value 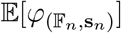 (readily computed via Eqn. (2.1)) when the number of species *n* is large. This condition allows some species to contribute proportionately more FD than other species do on average, and this proportion can grow with *n*, provided that it does not grow too quickly.

#### Proposition 2.3

a. For ϵ > 0,

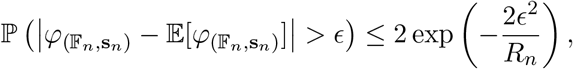

where 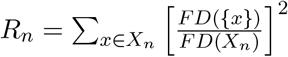.
b. If *R*_*n*_ *→* 0 as *n → ∞* then 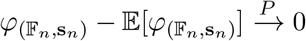.
c. Let av*FD*(*X*_*n*_) = *FD*(*X*_*n*_)*/n* (the average contribution of each species to the total FD), and suppose that for each *x* ∈ *X*_*n*_, *FD*({*x*})*/*av*FD*(*X*_*n*_) *≤ B*_*n*_, where and 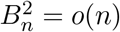 (e.g. 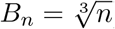). We then have the following convergence in probability as *n → ∞*:

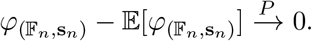

*Proof*. Let **Y**_**n**_ = {*Y*_*i*_ : *i ∈* [*n*]} be a sequence of Bernoulli random variables where each *Y*_*i*_ takes the value of 1 if species *x*_*i*_ survives and 0 otherwise. For the g-FOB model, the random variables *Y*_*i*_ are independent. We can write *φ*(𝔽_*n*_,**s**_*n*_) = *h*(**Y**_**n**_) where *h*(*y*_1_, …, *y*_*n*_) is the ratio 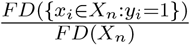. Observe that for any particular value of *i* ∈ {1, …, *n*}, if we change *y*_*i*_ (from 0 to 1 or visa versa) to give 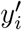 then:

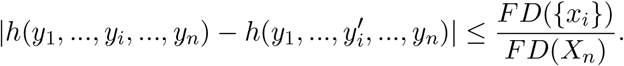

We now apply McDiarmid’s inequality [12] to obtain (for each ϵ > 0):

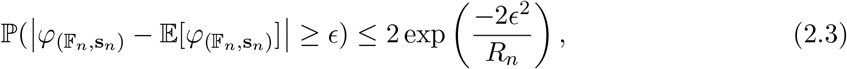

This establishes Part (a). Part (b) now follows immediately, and Part (c) follows from Part (b), since the condition in Part (c) implies that

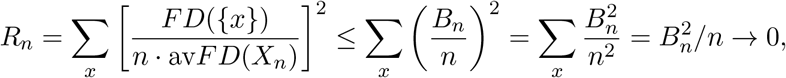

as *n → ∞*. □

**Remark**

Note that Proposition 2.3(c) can fail when the condition stated in Part (c) does not hold, even for the simpler FOB model. We provide a simple example to demonstrate this. Let *X*_*n*_ = {1, …, *n*} and let *F*_*i*_ = {*f*} for *i* = 1, …, *n −* 1 and *F*_*n*_ = {*g*} where *f* and *g* are distinct features with *μ*(*f*) = *μ*(*g*) = 1. In this case:

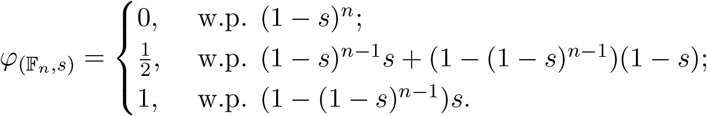

Therefore, 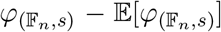 does not converge in probability to 0 (or to any constant) as *n → ∞*.

### 2.3 Consequences for Phylogenetic Diversity

Proposition 2.2 and Proposition 2.3 provide a simple way to derive certain results concerning phylogenetic diversity - both on rooted trees and also for rooted phylogenetic networks (specifically for the subNet diversity measure described in [24]). To each edge *e* of a rooted phylogenetic tree (or network), associate some unique feature *f*_*e*_ and give it the value *μ*(*f*_*e*_) = *𝓁*(*e*), where *𝓁*(*e*) is the length of edge *e* in the tree (or network). For any subset *Y* of *X* (the leaf set of *T*) we then have:

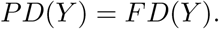

It follows that for the simple field of bullets models, PD satisfies the concavity properties described in Proposition 2.2, where the condition that the sets *F*_*x*_ are pairwise disjoint corresponds to the tree being a star tree. These results were established for rooted trees by specific tree-based arguments (see e.g. Section 5 of [8]), but they directly follow from the more general framework above, and extend beyond trees.

Using this same link between PD and FD, Proposition 2.3 provides a further application to any sequence of rooted phylogenetic trees *T*_*n*_ with *n* leaves and ultrametric edge lengths. For example, the ratio of surviving PD to original PD under the FOB model converges in probability to the expected value of this ratio for a sequence of trees *T*_*n*_ if the total PD of *T*_*n*_ grows at least as fast as *nL/n*^*β*^, where *L* is the height of the tree and 0 *< β <* 1*/*2. This condition holds, for example, for Yule trees [20].

## 3 The feature diversity ratio *φ* for a model of feature evolution on a phylogenetic tree

Consider a rooted binary phylogenetic tree *T*_*n*_, in which each edge has a positive length that corresponds to a temporal duration, with the root *ρ* of *T*_*n*_ being placed at the top of a stem edge at time 0, and with each leaf in the leaf set *X*_*n*_ = {*x*_1_, …, *x*_*n*_} of *T*_*n*_ being placed at time *t* (as in Fig. 3.1). For convenience, we will assume in this section that *μ*(*f*) = 1 for all features; however, this assumption can be relaxed (e.g. by allowing *μ*(*f*) to take values in a fixed interval [*a, b*] where *a >* 0 according to some fixed distribution, and independently between features) without altering the results significantly.

**Figure 3.1.**
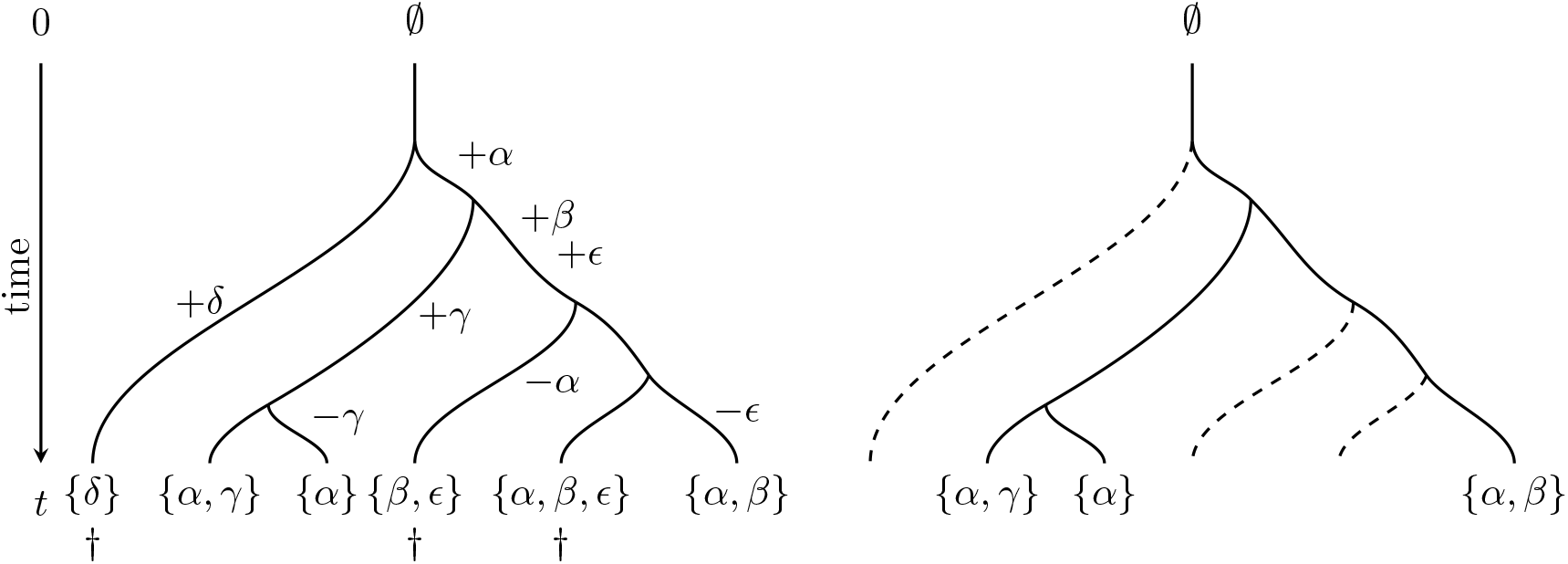
An example of feature gain and loss on a tree and the impact of extinction at the present. *Left:* New features arise (indicated by +) and disappear (indicated by *−*) along the branches of the tree. To simplify this example, no features are present at time 0; however, our results do not require this restriction. In total, there are five features present amongst the leaves of the tree at time *t*, namely {*α, β, γ, δ, ϵ*}. *Right:* An extinction event at the present (denoted by *†*) results in the loss of three of the extant species, leaving just three features being present among the leaves of the resulting pruned tree, namely {*α, β, γ*}. Thus, the ratio of surviving features to total features is 3*/*5 = 0.6.

We let *F*_*ρ*_ denote the (possibly empty) set of features present at time 0 (i.e. at the top of the stem edge), and we assume throughout this section that |*F*_*ρ*_| is bounded by some fixed constant *B*, independent of *n*.

On *T*_*n*_, we apply a stochastic process in which (discrete) features arise independently along the branches of this tree at rate *r*, and each feature that arises is novel (i.e. it has not appeared earlier elsewhere in the tree). Once a feature arises, it is then carried forward in time along the branches of *T*_*n*_ (and is passed on to the two lineages arising at any speciation event). In addition, any feature can be lost from a lineage at any point according to a continuous-time pure-death process that operates at rate *v*. This model was investigated in a different setting in [5] and studied more recently in [18]. Under this process, each leaf *x* of *T*_*n*_ will have a (possibly empty) set of features (*F*_*x*_). Fig. 3.1 illustrates the processes described. Note that |*F*_*x*_| (for any *x* ∈ *X*_*n*_) and *FD*(*X*_*n*_) are now random variables.

Let *N*_*𝓁*_ denote the number of features at the end of any path *P* in *T*_*n*_ that starts at time *t* = 0 and ends at time *𝓁*. Then *N*_*𝓁*_ is described by a continuous-time Markov process that has a constant birth rate and a linear death rate. It is then a classical result [3] that *N*_*𝓁*_ has expected value 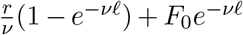, where *F*_0_ = |*F*_*ρ*_| (the number of features present at time 0), and *N*_*𝓁*_ converges to a Poisson distribution with mean 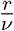 as *𝓁* grows. Moreover, if *N*_0_ = 0 then *N*_*𝓁*_ has a Poisson distribution for any value of *𝓁 >* 0 ([3] p. 461), and so, regardless of the value of *F*_*ρ*_, the random variable |*F*_*x*_ *\ F*_*ρ*_| has a Poisson distribution with mean 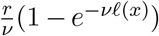 where *𝓁*(*x*) is the length of the path from the root to leaf *x*. The (random) number of features at any leaf *x* of *T*_*n*_ (i.e. |*F*_*x*_|) has the same distribution as *N*_*𝓁*(*x*)_. Note that for distinct leaves *x* and *y* of *T*_*n*_, the random variables |*F*_*x*_| and |*F*_*y*_| are not independent.

### Notational convention

Henceforth we will write *FD*(*T*_*n*_) in place of *FD*(*X*_*n*_) and we will also write 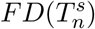 in place of *FD*(*χ*_*n*_) when *χ*_*n*_ is the subset of the set of leaves *X*_*n*_ of *T*_*n*_ that survive under a FOB model with a survival probability *s*. We will let *F*_*ρ*_ denote the set of features present at the root vertex *ρ* at the top of the stem edge.

#### Lemma 3.1

Set *F*_0_ = *∅*. Then for any value of *n ≥* 1 the following hold:

i. The random variable *FD*(*T*_*n*_) has a Poisson distribution.
ii. The expected value of *FD*(*T*_*n*_) satisfies the following bound:

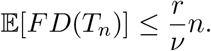

*Proof. Part* (*i*): We use induction on *n*. Since *F*_*ρ*_ = *∅*, the result for base case (*n* = 1) holds by the results mentioned in the previous paragraph. Thus, suppose that *n ≥* 2, and let *T*_*n*_ be a binary tree with *n* leaves and with *F*_*ρ*_ = *∅*. Then:

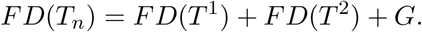

where the trees *T* ^*i*^ (with *i* = 1, 2) are subtrees of *T*_*n*_ obtained by deleting the stem edge and its endpoints (and setting the set of features at the top of the stem edge of *T* ^1^ and *T* ^2^ equal to the empty set), and *G* is the number of features that arise on the stem edge of *T*_*n*_ and are present in at least one leaf of *T*_*n*_.

Notice that *FD*(*T* ^1^), *FD*(*T* ^2^) and *G* are independent random variables, and, by induction, *FD*(*T* ^1^) and *FD*(*T* ^2^) each have a Poisson distribution. Conditional on the number *X* of features that are present at the end of the stem edge, *G* has a binomial distribution with parameters *X* and *p* where *p* is the probability that a single feature present at the end of the stem edge is present in at least one leaf of *T*_*n*_. Since *X* has a Poisson distribution, and a Poisson number of Bernoulli random variables is Poisson, it follows that *FD*(*T*_*n*_), being the sum of three independent Poisson variables, also has a Poisson distribution. This establishes the induction step and thus Part (i).

*Part* (*ii*): We have:

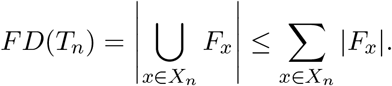

Now, for each *x* ∈ *X*, we have 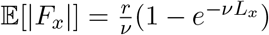 where *L*_*x*_ denotes the length of the path from the top of the stem edge of *T*_*n*_ to the leaf *x*. Thus 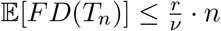. □

**Example (***n* = 2) Consider the process described on *T*_2_ with *F*_0_ = *∅*. Let *𝓁*_0_ denote the length of the stem edge, and *𝓁* the length of each of the two pendant edges. Then *FD*(*T*_2_) has a Poisson distribution with expected value

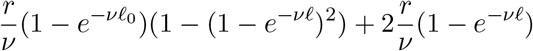

and 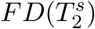 has expected value

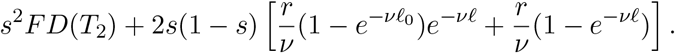

In particular,

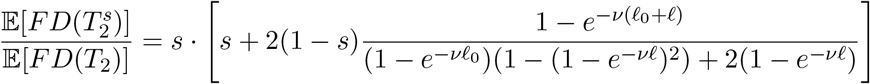

When *𝓁*_0_ = 0, the right-hand side of this equation equals *s*, but for all other values it is strictly greater than *s*. Moreover, by differentiating 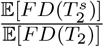 it can be verified that this ratio is monotone decreasing as *v* increases for all positive values of *𝓁* and *𝓁*_0_; in particular, for *s* ∈ (0, 1), and *ν >* 0, this ratio is always less than the expected proportion of PD that survives in *T*_2_.

### 3.1 A limit result for sequences of trees

The main result of this section is Theorem 3.1, and its proof relies on establishing a sequence of preliminary lemmas.

#### Lemma 3.2

Let *β >* 0 be a fixed constant. Given a sequence *T*_*n*_ of trees with leaf set *X*_*n*_, let *ε*_*n*_ be the event that |*F*_*x*_| *≤ n*^*β*^ for every *x* ∈ *X*_*n*_. Then lim_*n→∞*_ ℙ(*ε*_*n*_) = 1.

*Proof*. We combine the Bonferroni inequality with a standard right-tail probability bound for a Poisson variable. Firstly, observe that:

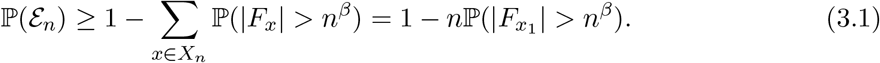

Now, 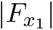 can be written as the sum of two independent random variables 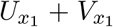 where 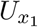 has a Poisson distribution with mean *m ≤ r/v*, and 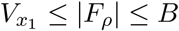 with probability 1 (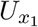 counts the features at *x*_1_ if 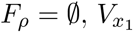 counts the features in *F*_*ρ*_ that are remain present at *x*_1_, and *B* is the global bound on *F*_*ρ*_ described near the start of Section 3). Thus, the Chernoff bound on the right hand tail of a Poisson variable ([14], p. 97) gives 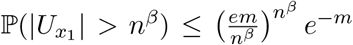, and so *n*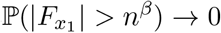 as *n* grows. Applying this to the inequality in (3.1) establishes the result.

#### Lemma 3.3

Let (*T*_*n*_, *n ≥* 1) be a sequence of rooted binary trees, with *T*_*n*_ having leaf set *X*_*n*_, and suppose that 𝔼[*FD*(*T*_*n*_)] *≥ cn* for some constant *c >* 0. Then 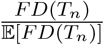 converges in probability to 1 as *n → ∞*.

*Proof*. First observe that we can write *FD*(*T*_*n*_) as the sum of two independent random variables, namely *FD*_0_(*T*_*n*_) + *K*, where *FD*_0_(*T*_*n*_) is the *FD* value when *F*_*ρ*_ = ∅, and *K* is the number of features at the time at the root that are also present in at least one leaf. In particular, *K ≤* | *F*_*ρ*_ | which is assumed to be bounded by a constant *B* (independent of *n*). Thus

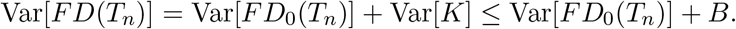

By Lemma 3.1, 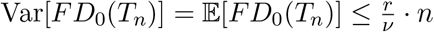, so 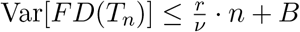, and thus:

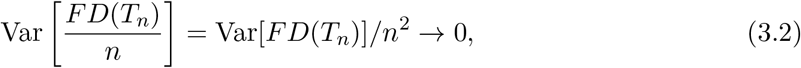

as *n → ∞*. We now apply Chebyshev’s inequality to obtain:

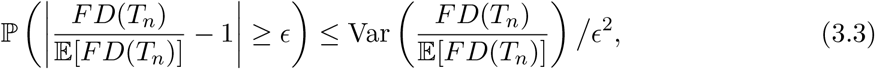

and since

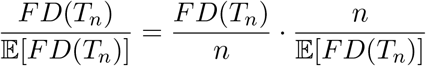

we have:

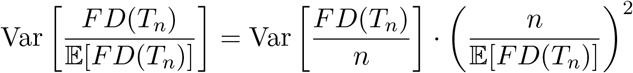

and so the term on the right of Eqn. (3.3) is bounded above by 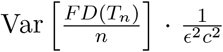, which converges to 0 as *n → ∞* by Eqn. (3.2).

□

#### Lemma 3.4

Let (*T*_*n*_, *n ≥* 1) be a sequence of rooted binary trees, with *T*_*n*_ having leaf set *X*_*n*_, and suppose that 𝔼[*FD*(*T*_*n*_)] *≥ cn* for some constant *c >* 0. Suppose that for each *x* ∈ *X*_*n*_, *FD*({*x*}) ≤ *B*_*n*_, where 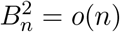. Then 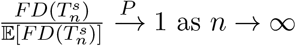:

*Proof*. Let *W*_*n*_ = *H*(**Y**_**n**_), where **Y**_**n**_ = {*Y*_*i*_ : *i ∈* [*n*]} is the sequence of Bernoulli random variables with *Y*_*i*_ = 1 if species *x*_*i*_ survives the FOB extinction event and *Y*_*i*_ = 0 otherwise, and *H*(*y*_1_, …, *y*_*n*_) is the ratio 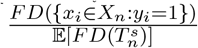, where the numerator is as defined in Eqn. (2.1) with *μ*(*f*) = 1 for all *f*. Observe that for any particular value of *i* ∈ {1, …, *n*}, if we change *y*_*i*_ (from 0 to 1 or visa versa) to give 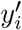 then:

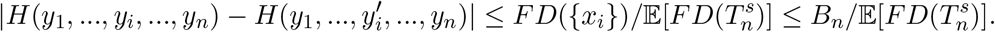

We now apply McDiarmid’s inequality to obtain (for each *ϵ >* 0):

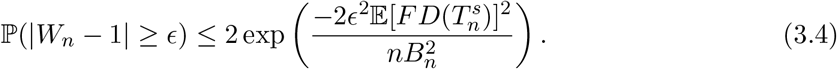

Now, 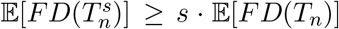 by Proposition 2.2(i) (taking *μ*(*f*) = 1 for all *f*) and, by assumption, 𝔼[*FD*(*T*_*n*_)] *≥ cn*. Thus we obtain:

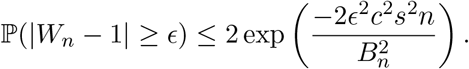

Therefore, ℙ(|*W*_*n*_ *−* 1| *≥ ϵ*) *→* 0 as *n → ∞*. Since this holds for all ϵ *>* 0, we obtain the claimed result.

□

We can now state the main result of this section. Recall that 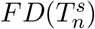 is the number of features present among leaves of *T*_*n*_ that survive the FOB extinction event.

#### Theorem 3.1

*Let* (*T*_*n*_, *n ≥* 1) *be a sequence of binary trees and let features evolve on T*_*n*_ *according to the stochastic feature evolution process described. If* 𝔼[*FD*(*T*_*n*_)] *≥ cn for some constant c >* 0, *and if* 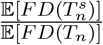 *converges to a constant c*_*s*_ *as n grows we have:*

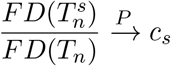

*as n → ∞*.

Before we proceed to the proof, we provide the following comments.

**Remarks**

- Theorem 3.1 can fail without the condition 𝔼[*FD*(*T*_*n*_)] *≥ cn*. Fig. 3.2 provides a simple example of a sequence of trees *T*_*n*_ for which 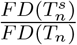 does not converge in probability to any constant value as *n* grows. In this example, *F*_*ρ*_ = *∅* and the tree has height 2*𝓁* with one leaf having an incident edge of length *𝓁* and the remaining *n−* 1 leaves having incident edges of length 1*/n*.

**Figure 3.2.**
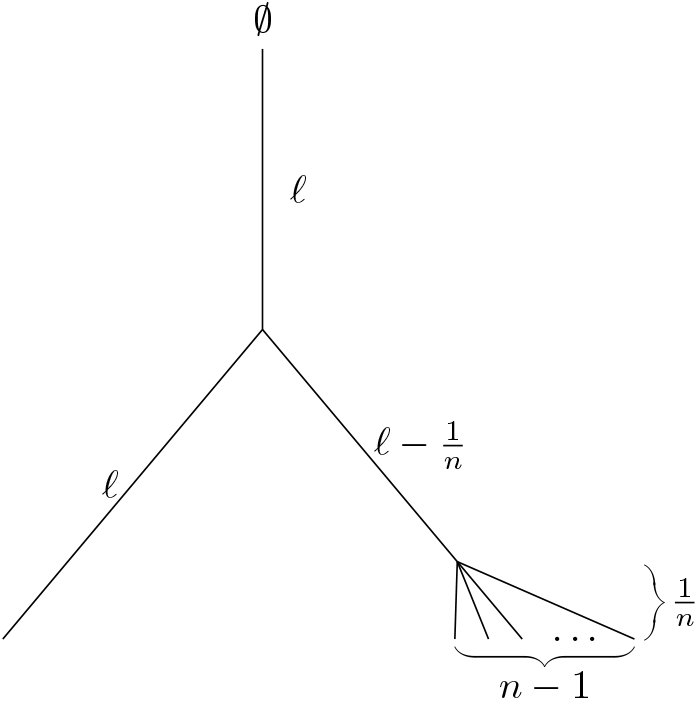
For the tree *T*_*n*_ shown (with *𝓁>* 1 fixed and *s* ∈ (0, 1)) the sequence of trees (*T*_*n*_, *n ≥* 2) has the property that 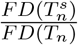 does not converge in probability to any constant value as *n* grows.
- A sufficient condition for the inequality 𝔼 [*FD*(*T*_*n*_)] *≥ cn* in Theorem 3.1 to hold is that for some *ϵ >* 0 and *δ >* 0 the proportion of pendant edges of *T*_*n*_ of length *≥ ϵ* is at least *δ* for all *n ≥* 1. Briefly, the reason for this is that the expected number of features arising on each of these pendant edges and surviving to the end of this edge is bounded away from 0 and the features associated with distinct pendant edges are always different from each other, and different from any other features arising in the tree.

*Proof of* Theorem 3.1.

Let *A* _*ϵ*_ (*n*) be the event that 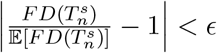, and let *ε*_*n*_ be the event described in Lemma 3.2 with 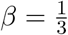. Then, lim_*n→∞*_ ℙ(*A*_*ϵ*_ (*n*)| *ε*_*n*_) = 1, by Lemma 3.4. Now,

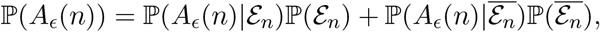

and since ℙ(*ε*_*n*_) *→* 1 as *n → ∞* (by Lemma 3.2) we have lim_*n→∞*_ ℙ(*A* _*∈*_ (*n*)) = 1. Since this holds for all *ϵ >* 0, it follows that 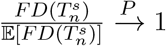 as *n* grows.

Moreover, from Lemma 3.3, we have 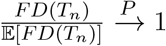. Now we can write 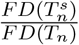 as follows:

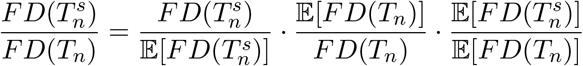

The first two terms on the right of this equation each converge in probability to 1 as *n → ∞*, whereas the third (deterministic) term converges to *c*_*s*_. This completes the proof. □

We will apply Theorem 3.1 in the next section to establish a result for FD loss on birth-death trees.

## 4 Feature diversity ratios in birth–death trees

In this section, we continue to investigate the stochastic model of feature gain and loss, but rather than considering fixed trees, we will now allow the trees themselves to be stochastically generated, following the simple birth–death processes that are common in phylogenetics. Thus, there will now be *three* stochastic processes in play: the linear-birth/linear-death process that generates the tree, the constant-birth/linear-death process of feature gain and loss operating along the branches of the tree, and the simple FOB extinction event at the present.

### 4.1 Definitions

Let *𝒯*_*t*_ denote a birth–death tree grown from a single lineage for time *t* with birth and death parameters *λ* and *μ*, respectively. We will assume throughout that *λ > μ* (since otherwise the tree is guaranteed to die out as *t* becomes large).

On *𝒯*_*t*_, we impose the model of feature gain and loss from the previous section with parameters *r* and *v*. We now apply the FOB model in which each extant species (i.e. leaves of *𝒯*_*t*_ that are present at time *t*) has a probability *s >* 0 of surviving and 1 *− s* of becoming extinct (independently across the extant species), and we let 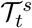 denote the tree obtained from *𝒯*_*t*_ by removing the species at the present that did not survive this process. We refer to 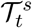 as the *pruned* tree, and the leaves of *𝒯*_*t*_ and 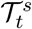 that are present at time *t* as the *extant species* (*or leaves*) of these trees (to contrast them from leaves of *𝒯*_*t*_ that lie at the endpoints of any extinct lineages). If *s <* 1, there may now be fewer features present among the (probably reduced number of) extant species in 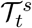 than there were in *𝒯*_*t*_.

Let *FD*(*𝒯*_*t*_) be the discrete random variable that counts the number of features present in at least one of the extant leaves of *𝒯*_*t*_, and let 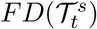 denote the number of these features that are also present in at least one extant leaf in the pruned tree 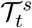.

### 4.2 Expected feature diversity

Next, we consider the expected values of *FD*(*𝒯*_*t*_) and 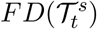, where this expectation is across all three processes (the birth–death process that generates *𝒯*_*t*_, feature gain and loss on this tree, and the species that survive the extinction event at the present under the FOB model). Of particular interest is the ratio of these expectations, and their limit as *t* becomes large. Specifically, let:

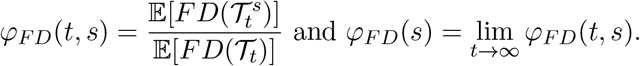

Note that *φ*_*FD*_(*s*) is a function of five parameters (*s, r, λ, μ, v*); however, we will show that it is just a function of *s* and two other parameters. Notice also that once these parameters are fixed, *φ*_*FD*_ (*t, s*) and *φ*_*FD*_ (*s*) are monotone increasing functions of *s* taking the value 0 at *s* = 0 and 1 at *s* = 1. Moreover, *ϕ*_*F D*_(*t, s*) and *ϕ*_*FD*_ (*s*) are both independent of *r* (the rate at which features arise along any lineage) as we formally show shortly.

### 4.3 Relationship to Phylogenetic Diversity (PD)

Recall that for a rooted phylogenetic tree *T* with branch lengths, the PD value of a subset *S* of leaves (*PD*(*S, T*)) is the sum of the lengths of the edges of the subtree of *T* that connect *S* and the root of the tree.

In the special case where *v* = 0, and where no features are present at time 0, 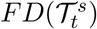 (conditioned on *𝒯*_*t*_), has a Poisson distribution with a mean of *r* times *PD*(*𝒮*_*t*_, *𝒯*_*t*_), where *𝒮*_*t*_ is the (random) set of leaves at time *t* that survive the extinction event at the present. Consequently, 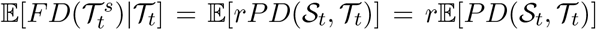. Similarly, *𝔼* [*FD*(*𝒯*_*t*_)| *𝒯*_*t*_] = *r 𝔼* [*PD*(*ℒ*_*t*_, *𝒯*_*t*_)], where *ℒ*_*t*_ is the set of extant leaves of *𝒯*_*t*_. Thus, in this special case we have *φ*_*FD*_ (*s*) = *φ*_*PD*_ (*s*), where:

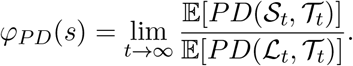

The function *φ*_*PD*_ (*s*) was explicitly determined in [8, 15] as follows:

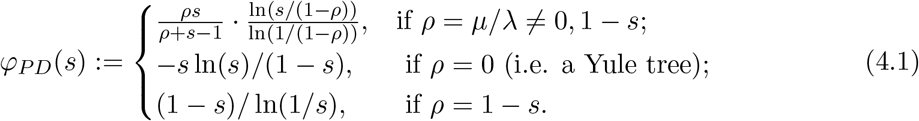

## 5 Calculating *φ*_*FD*_(*s*)

We first recall a standard result from birth–death theory. Consider a linear birth–death process (starting with a single individual at time 0), with a birth rate *λ*, a death rate *θ*. For the individuals present at time *t*, sample each individual independently with sampling probability *s >* 0. Let *X*_*t*_ (*t ≥* 0) denote the number of these sampled individuals and let 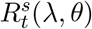 be the probability that *X*_*t*_ *>* 0. Then

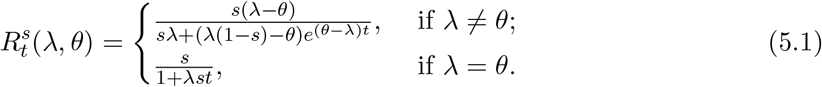

In particular, 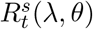 converges to 0 if *λ ≤ θ* and converges to a strictly positive value 1 *− θ/λ* if *λ> θ* [7, 26].

The number of species at time *t* in 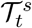 that have a copy of particular feature *f* that arose at some fixed time *t*_0_ ∈ (0, *t*) in *𝒯*_*t*_ is described exactly by the birth–death process 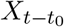 with parameters *λ* and *θ* = *μ*+*v* and survival probability *s* at the present; it follows that as *t* becomes large, it becomes increasingly certain that none of the species at time *t* in the pruned tree will contain feature *f* if *λ ≤ μ* + *v*, whereas if *λ > μ* + *v*, there is a positive limiting probability that *f* will be present in the extant leaves of the pruned tree.

Since *φ*_*FD*_ (*s*) = *φ*_*PD*_ (*s*) (as given by Eqn. (4.1)) for all values of *s* when *v* = 0, in this section we will assume that *v >* 0 (in addition to our universal assumption that *λ > μ*). Our main result provides an explicit formula for *φ*_*FD*_ (*s*) in Part (a), and describes some of its key properties in Parts (b) and (c).

### Theorem 5.1

*Given λ > μ and v >* 0, *let* 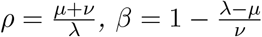. *Then:*

a. 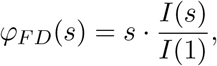

*where*

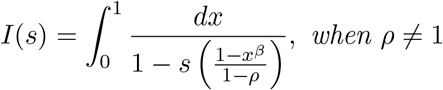

*and*

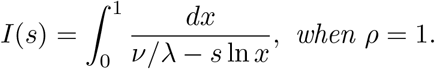
b. *Conditional on the non-extinction of T*_*t*_, 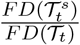 *converges in probability to φ*_*FD*_ (*s*) *as t → ∞*.
c. *φ*_*FD*_ (*s*) *is an increasing concave function that satisfies* 1 *≥φ*_*FD*_ (*s*) *≥ s for all s*.

**Remarks**

i. Notice that although *φ*_*FD*_ (*s*) depends on five parameters (*r, s, λ, μ, v*), Theorem 5.1(a) reveals that just three derived parameters suffice to determine *ϕ*_*FD*_ (*s*), namely *s* and the ratios *ρ*_1_ = *μ/λ* ∈ (0, 1) and *ρ*_2_ = *v/λ* (these determine *ρ* and *β*, since *ρ* = *ρ*_1_ + *ρ*_2_ and *β* = 1 *−* 1*/ρ*_2_ + *ρ*_1_*/ρ*_2_). Notice also that *ρ* = 1 *⇔ β* = 0 and *ρ >* 1 *⇔ β >* 0.
ii. Our proof relies on establishing the following exact expression for 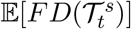:

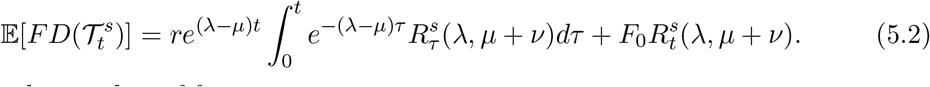

Where *F*_0_ is the number of features present at time *t* = 0. In particular, this also provides an exact expression for *ϕ*_*F D*_(*t, s*). Notice that the ratio of the expected number of features present in the pruned tree, divided by the expected number of species in the pruned tree is 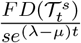, and this ratio converges to 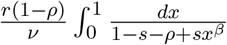 as *t → ∞* when *ρ* ≠ 1 (via a further analysis of Eqn. (5.2) in the Appendix).
iii. We saw in Section 4.3 that when *v* = 0, *φ*_*FD*_ (*s*) = *φ*_*PD*_(*s*). At the other extreme, if *λ* and *μ* are fixed, and we let *v → ∞*, then *φ*_*FD*_ (*s*) converges to *s* (since *ρ → −∞* and *β →* 1 in Theorem 5.1(a)). Informally, when *v* is large compared to *λ*, most of the features present among the extant leaves of *𝒯*_*t*_ will have arisen near the end of the pendant edges incident with these extant leaves (a formalisation of this claim appears in [18]); if we now apply the FOB model then the expected proportion of these features that survive will be close to the expected proportion of leaves that survive, namely *s*.

## 5.1 Illustrative examples

First, consider a Yule tree (i.e. *μ* = 0) grown for time *t*. Figure 5.1 (left) plots *φ*_*FD*_ (*s*) for values of *v/λ ∈* {0, 0.5, 1, 2, 10}. When *v* = 0, *φ*_*FD*_ (*s*) describes the proportional loss of expected PD in the pruned tree, and as *v* increases, *φ*_*FD*_ (*s*) converges towards *s* (the expected proportion of leaves that survive extinction at the present). Figure 5.1 (right) plots *φ*_*FD*_ (*s*) for birth–death trees with *μ/λ* = 0.8, showing a similar trend, however with *φ*_*FD*_ (*s*) ranging higher above the curve *y* = *s*.

**Figure 5.1.**
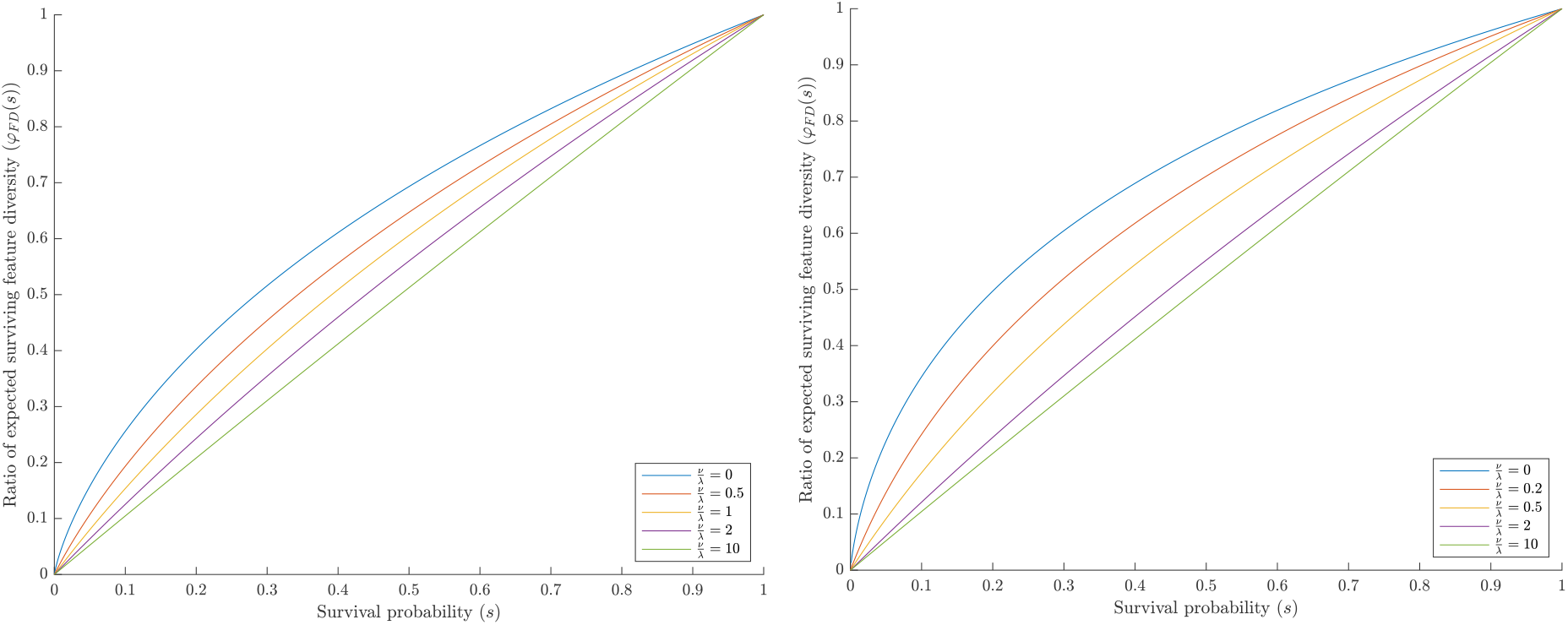
*Left:* The function *φ*_*FD*_ (*s*) as a function of *s* for pure-birth Yule trees (*μ* = 0). *Right:* The function *φ*_*FD*_ (*s*) for birth–death trees with *μ/λ* = 0.8. The curves within each graph show the effect of increasing *v*, for values of *v/λ* between 0 (top) and 10 (bottom). The top curve also corresponds to the phylogenetic diversity ratio *φ*_*PD*_ (*s*).

## 5.2 Simulations

We ran simulations to test the expected relationship between *φ*_*FD*_ (*s*) and *s* and to get estimates of standard deviation. All simulations were run in R version 4.2.1 [21]. We first simulated 500 Yule trees (age = 100, *λ* = 0.055, *μ* = 0, repeated and filtered to keep 500 250-tip trees) and 500 birth-death trees (age = 100, *λ* = 0.11, *μ* = 0.088, repeated and filtered to keep 500 trees with 250-300 tips) using the sim.bd.age function in the package TreeSim [19]. Features were then evolved on each tree, followed by separate extinction events.

Keeping the rate of feature gain fixed (*r* = 0.3), we modelled five different rates of feature loss (*v ∈* {0, 0.5*λ, λ*, 2*λ*, 10*λ*}). We estimated feature gain and loss on each edge using the Gillespie algorithm [4], where time until the next event (either feature gain or loss) was drawn from an exponential distribution with rate (*r* + *kv*), where *k* is the number of currently existing features at the start of the edge. The type of event was then determined with a Bernoulli draw with probability of a gain equal to *r/*(*r* + *kv*). At each split on the tree, all existing features were copied to descendent edges. Each gain event created a new unique feature, and each loss event randomly selected an existing feature to eliminate from the current edge. At the end of the simulation the presence of features on each tip of the tree was recorded.

Extinction events were simulated by randomly selecting a proportion *s* of tips to delete, with *s* ranging from 0.05 to 0.95 by intervals of 0.05. The proportion of unique features remaining after extinction events was recorded, and the results are shown in Fig. 5.2.

**Figure 5.2.**
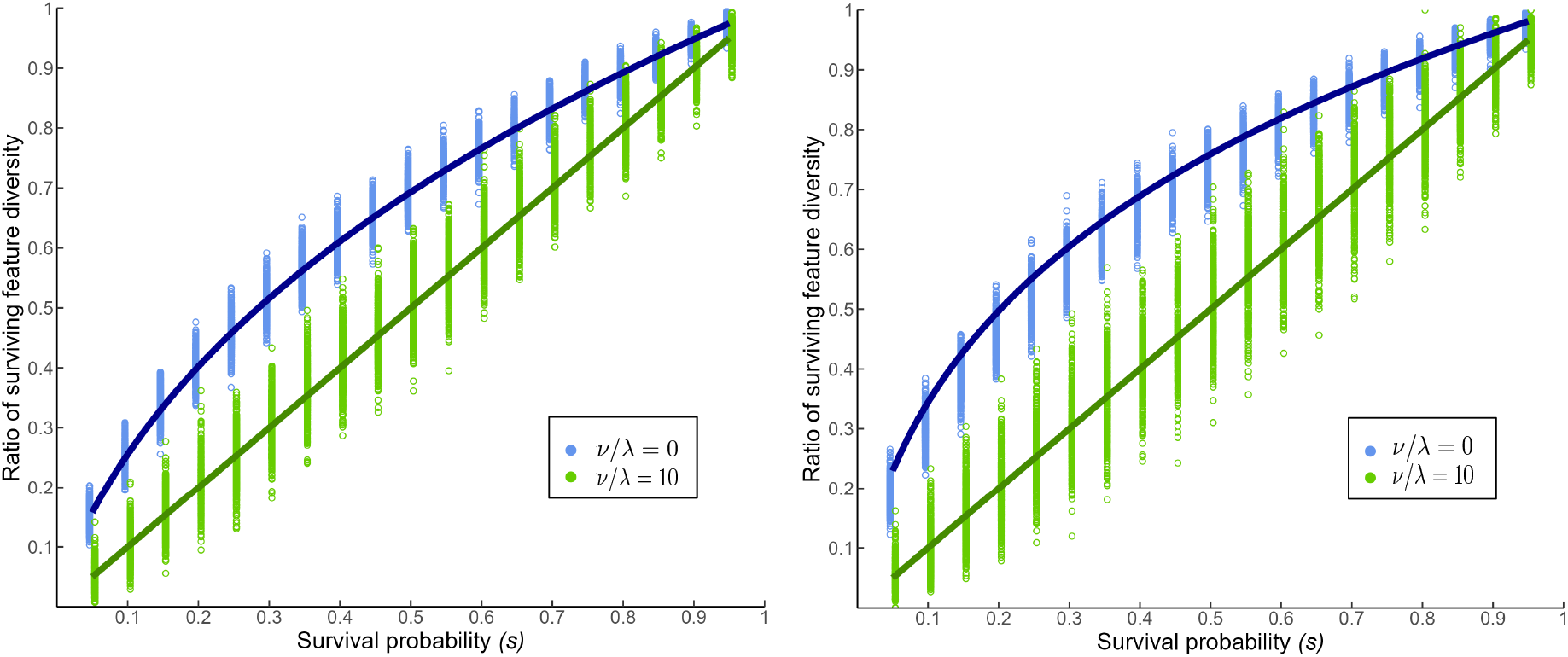
Simulation results for remaining feature diversity as a function of *s* for pure-birth Yule trees (*μ* = 0, left) and birth-death trees (*μ/λ* = 0.8, right). Results shown for two of the five feature evolution parameter scenarios. Each point represents one simulation on one simulated tree (*n* = 500 trees). The solid curves are the theoretical relationships between the expected proportion of remaining feature diversity and *s* for both of the shown feature evolution parameter scenarios.

Resulting ratios of remaining features generally tracked expectations (the bias for birth-death trees when *v/λ* = 0 is likely due to our theoretical results conditioning on *t* rather than *n*). The standard deviation (SD), calculated for each *v* on both Yule and birth-death trees, were fairly consistent for all *v* except *v* = 10*λ*, where it was noticeably higher, as shown in Table 1. Because the number of total events along an edge (gains and losses) is described by a Poisson distribution, its variance increases with the mean, and this may explain the higher standard deviation at the highest loss rate.

**Table 1.**
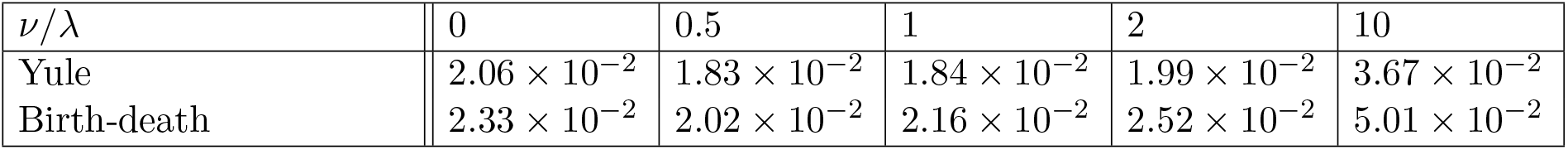
Mean standard deviations of proportions of remaining feature diversity (each value is averaged over the 19 values of *s* between 0.05 and 0.95).

| $\nu/\lambda$ | 0 | 0.5 | 1 | 2 | 10 |
| --- | --- | --- | --- | --- | --- |
| Yule | $2.06 \times 10^{-2}$ | $1.83 \times 10^{-2}$ | $1.84 \times 10^{-2}$ | $1.99 \times 10^{-2}$ | $3.67 \times 10^{-2}$ |
| Birth-death | $2.33 \times 10^{-2}$ | $2.02 \times 10^{-2}$ | $2.16 \times 10^{-2}$ | $2.52 \times 10^{-2}$ | $5.01 \times 10^{-2}$ |

## 5.3 Features that appear in only one extant species

Let *𝒰*_*t*_ denote the number of features that are present in precisely one species in *𝒯*_*t*_, and let *U*_*t*_ = 𝔼[*𝒰*_*t*_]. The following result describes a simple relationship between *U*_*t*_ and *F*_*t*_ = 𝔼[*FD*(*𝒯*_*t*_)].

### Proposition 5.1

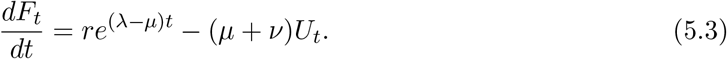

*Proof*. Let *ℱ*_*t*_ denote the number of features present at time *t* among the leaves of *𝒯*_*t*_. Consider evolving *𝒯*_*t*_ for an additional (short) period *δ* into the future. Then, conditional on *𝒩*_*t*_ (the number of leaves of *𝒯*_*t*_ present at time *t*) and *𝒰*_*t*_:

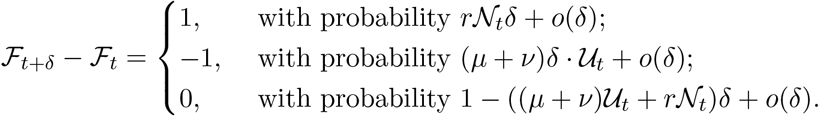

Applying the expectation operator (and using 𝔼[*ℱ*_*t*+*δ*_] = 𝔼 [𝔼[*ℱ*_*t*+*δ*_|*𝒩*_*t*_, *𝒰*_*t*_]]) and letting *δ →* 0 leads to the equation stated.

□

## 6 Concluding comments

In this paper, we have considered, in order, three types of data to quantify the expected loss of feature diversity: sets of features across species (without any model of feature evolution or phylogeny), sets of features at the tips of a given phylogenetic tree, and sets of features at the tips of a random (birth–death) tree. The results of the earlier sections also proved helpful in establishing certain results in later sections.

In terms of wider significance to biodiversity conservation, our results and graphs in Section 5 suggest that the extent of relative feature diversity loss following extinction at the present is likely to be greater than that predicted by relative phylogenetic diversity loss for any given extinction rate *s* ∈ (0, 1).

Of course, our results are based on simple models (of feature gain and loss, and extinction at the present) and so exploring how these results might extend to more complex and realistic biological models would be a worthwhile topic for future work.

## Acknowledgements

We thank François Bienvenu and James Rosindell for helpful suggestions on an earlier draft of this manuscript, and Ailene MacPherson for technical advice regarding the simulations. We also thank the two anonymous reviewers for further helpful comments and the New Zealand Marsden Fund (MFP-UOC2005) for supporting this research.

## 8 Appendix: Proof of Theorem 5.1

### Part (a)

Consider a birth–death tree *𝒯*_*t*_ with parameters *λ, μ*, a feature evolution model with parameters *r, v*, and a survival probability *s* for leaves at the present. Let 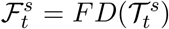, and let 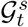 be the random variable that has the same distribution as 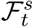 with the initial condition 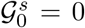 (i.e. no features at the root of the tree). Let 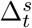 be the number of features that are present at the root of *𝒯*_*t*_ and also in the pruned tree 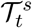. Then

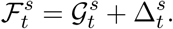

Let 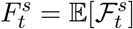 and 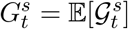. The two random variables 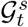 and 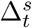 are not independent (they are linked by both the underlying tree *𝒯*_*t*_ and the pruning event at the present) however, applying the expectation operator gives:

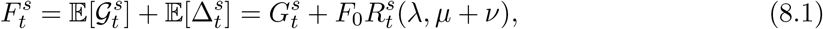

where *F*_0_ is the number of features present at the root of 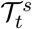.

Now, consider 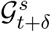, and the events that can occur in the interval [0, *δ*).

a. A new feature arises (with probability *rδ* + *o*(*δ*));
b. A speciation event occurs (with probability *λδ* + *o*(*δ*));
c. The lineage (and hence the tree) dies (with probability *μδ* + *o*(*δ*)).
d. None of the above occur (with probability 1 *−* (*r* + *λ* + *μ*)*δ* + *o*(*δ*))

Let *X* be the random variable taking values in {*a, b, c, d*} which denotes which of these four events occurs. In Case (a), the new feature is also present in 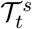 with probability 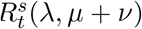 and so

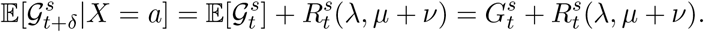

For Case (b), 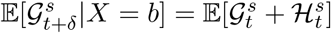, where 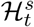 is an independent copy of 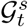. Thus,

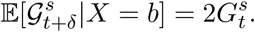

For Cases (c) and (d), we have: 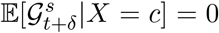 and 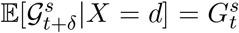. Thus, by the law of total expectation,

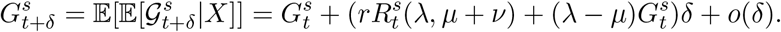

Consequently, the function 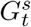 satisfies the first-order linear differential equation:

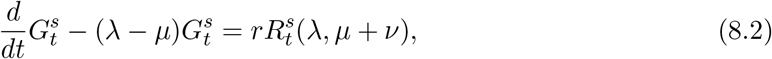

subject to the initial condition 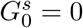. Solving Eqn. (8.2) gives:

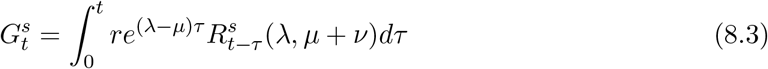

By making a change of variable we can rewrite Eqn. (8.3) as:

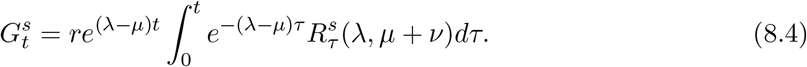

Combining this equation and Eqn. (8.1) provides the explicit expression for 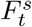 described earlier (Eqn. (5.2)).

We now substitute in the expression for 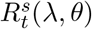 from Eqn. (5.1) (with *t* = *𝒯*and *θ* = *μ* + *v*). For *ρ≠* 1, we have:

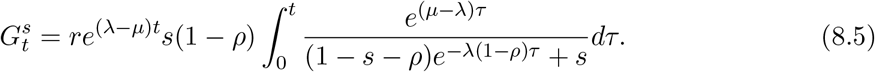

Multiplying the numerator and denominator of the integrand by *e*^*λ*(1*−ρ*) *𝒯*^ gives 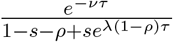 and then by making the substitution *x* = *e*^*−v 𝒯*^, we obtain:

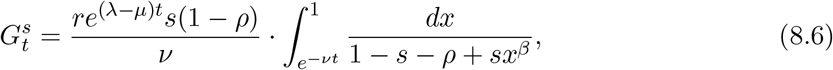

for the value of *β* described in the theorem. Combining Eqns. (8.1) and (8.6) gives:

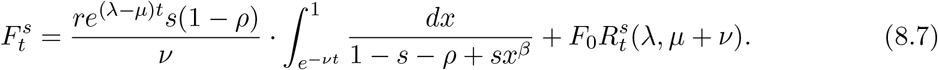

Thus, for *ρ≠* 1,

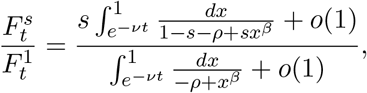

where the two terms of order *o*(1) (which converge to zero as *t* grows) refer to the last term on the right of Eqn. (8.7) which is bounded above by the constant *F*_0_ and so is asymptotically negligible in comparison to the term *e*^(*λ−μ*)*t*^ in Eqn. (8.6). This gives the limit for *φ*_*FD*_ (*s*) as stated in Part (a) for fixed *ρ≠* 1.

In the case where *ρ* = 1 (which implies *β* = 0), we use the corresponding expression for 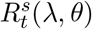 from Eqn. (5.1) with *t* = *𝒯*and *θ* = *μ* + *v* to obtain:

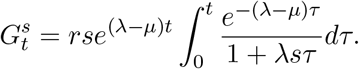

By a similar approach to the above we are led to the second equation in Part (a).

**Part (b)**

Let *𝒯*_*t*_ be a birth–death tree with rates *λ, μ* where *λ > μ*, let *n*(*𝒯*_*t*_) denote the number of leaves of *𝒯*_*t*_ present at time *t*, and let *ε*^*′*^ be the event that *n*(*𝒯*_*t*_) *>* 0 (i.e. the non-extinction of *𝒯*_*t*_).

Conditional on the event *ε*^*′*^, the number of leaves in *𝒯*_*t*_ tends to infinity (with probability 1) as *t → ∞* [6], and so we can define a sequence of trees *T*_1_, *T*_2_, …, *T*_*n*_, … by letting *T*_*k*_ denote the tree *𝒯*_*𝒯*_ at the first time *𝒯*= *𝒯*(*k*) when _*τ*_ has *k* extant leaves (we ignore leaves of _*t*_ that have already become extinct by time *τ*).

Next, we establish that 𝔼[*FD*(*T*_*n*_)] *≥cn* for a constant *c >* 0 (in order to apply Theorem 3.1). The tree *T*_*n*_ has *n* extant pendant edges, and the length of a randomly selected pendant edge in *T*_*n*_ has a strictly positive probability *p* of having length at least *κ >* 0 (dependent on *μ* and *λ*), by Theorem 3.1 of [20]. Now, 𝔼[*FD*(*T*_*n*_)] is bounded below by the total number of features that arises on the *n* pendant edges and survive to the end of the edge (since all these features will necessarily be distinct from each other, and from other features that arise in the tree). Moreover, for each edge having length at least *κ* the expected number of features that arise on this edge and survive to the end of the edge is at least 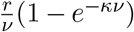. Thus the expected number of features contributed by the pendant edges to *FD*(*T*_*n*_) is at least 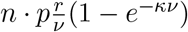.

Thus, we can now apply Theorem 3.1, since 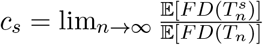 exists, and equals *φ*_*FD*_ (*s*), so 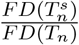 (and thus 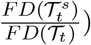 converges to *φ*_*FD*_ (*s*) as *n* (respectively *t*) grows.

**Part (c)**: We apply Proposition 2.2. By Part (i) of that result, and conditioning on *𝒯*_*t*_ we obtain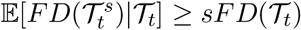, and so, taking expectation again (over the distribution of *𝒯*_*t*_) gives: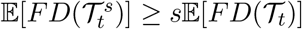, and thus 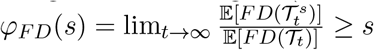.

The inequality *φ*_*FD*_ (*s*) *≤* 1 is clear since, for any choice of *𝒯*_*t*_, we have 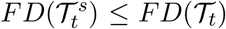 with probability 1.

For concavity, Proposition 2.2 implies that the conditional expectation 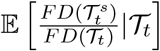 is concave as a function of *s*, and (by taking expectation over the distribution of *𝒯*_*t*_), 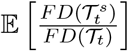 is also concave as a function of *s*. Finally, by Part (a) of the current theorem, 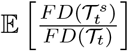 converges (deterministically) to *φ*_*FD*_ (*s*) and so this function is also concave as a function of *s*.

□

## Notes

### Competing Interest Statement

The authors have declared no competing interest.

### Summary of Updates

This version includes simulations (carried out by the additional author Daniel Pelletier) and corrects some typos in an earlier version of the paper

